# Benzidine Induces Proliferation of UMUC3 Bladder Cancer Cells Through Activation of ERK5 Pathway

**DOI:** 10.1101/2022.06.11.495720

**Authors:** Xin Sun, Tao Zhang, Liang Tang, Jie Min, Yi Wang, De-Xin Yu

**Affiliations:** Department of Urology, the Second Hospital of Anhui Medical University, Hefei, Anhui 230032, P.R. China

**Keywords:** benzidine, bladder cancer, cell proliferation, ERK5 pathway

## Abstract

Mounting evidences have revealed that benzidine, a known human carcinogen, is a pivotal risk factor for bladder cancer (BC). Dysregulated cell proliferation serves as a pivotal pathophysiological step in BC development. The potential mechanisms of MAPK pathways, particularly extracellular regulated protein kinases 5 (ERK5), regarding the modulation of benzidine-elicited cell proliferation are still elusive. Herein, human BC cells UMUC-3 were exposed to indicated levels of benzidine for 5 days. Cell growth, cell cycle progression, cell cycle-related, MAPK genetic expression, and related protein contents were investigated by the 3-(4,5-dimethylthiazol-2-yl)-2,5-diphenyltetrazolium bromide (MTT) test, flow cytometry, quantitative reverse transcription-polymerase chain reaction (qRT-PCR), and western blotting (WB), respectively. XMD8-92 (a specific suppressor of ERK5) and small interfering RNA (siRNA) were utilized to identify the roles of ERK5. Our results illustrated that benzidine elevated the proliferative ability of UMUC3 cells and accelerated the cell cycle transition from G1 to S phase, and it increased the expressions of cyclin D1 and proliferating cell nuclear antigen (PCNA) and reduced the expression of p21. Meanwhile, benzidine treatment resulted in the activation of ERK5 pathway and activator protein 1 (AP-1) proteins. In addition, benzidine-elicited cell proliferation and ERK5 stimulation were fully inhibited by XMD8-92 and siRNAs specific to ERK5. Taken together, these results demonstrated that ERK5-mediated cell proliferation was tightly related to benzidine-assoicated BC development, which established the foundation for targeting the ERK5 pathway in BC treatment.

## INTRODUCTION

Urinary BC is one of commonly diagnosed malignancies worldwide, with a yearly incidence of approximately 549,000 cases in 2018, which took up 3% of the entire newly diagnosed tumor cases [1]. Recently, the incidence and mortality of bladder cancer have been rising gradually. In 2022, the estimated number of new patients and mortalities in the US are 81180 and 17100, respectively [2]. Benzidine was predominantly utilized in the stain industry by makers of rubber, leather, textiles and paint products along with printing corporations [3]. Labelled as ‘the most important carcinogenic aromatic amine directed towards the bladder’, benzidine was categorized as definite mankind carcinogenic agents (Group 1) by the International Agency for Research on Cancer (IARC) in 1980 [4]. To date, extensive evidences indicate the positive association between benzidine and BC [3–5]. Nonetheless, the molecule-level etiopathogenesis is still elusive.

The cell cycle contains a range of tightly controlled events which result in the replication of genomic DNA and the production of 2 daughter cells [6]. In principle, cell cycle includes 4 stages: G1, S, G2, and M. Unless the previous phase is completed successfully, cells are banned to progress from one phase to another. The ability to sustain unscheduled proliferation is considered as a hallmark of cancer [7]. Mounting proofs have revealed that cell cycle deregulation is closely related to the developmental process of BC [8]. Additionally, it has been discovered that the regulatory role of cell cycle is integrated into complex molecular mechanisms, including the ERK5 pathway [9].

The MAPK pathway is composed of four signaling subfamilies: ERK1/2, the Jun N-terminal kinases (JNKs), p38, and ERK5. Multiple studies have revealed that the MAPK pathway participates in various kinds of cellular responses induced by environmental and developmental signal transduction [10, 11]. Amongst the MAPK family, ERK5 is a unique kinase, and it has a large structure featured by the existence of a special C-terminal domain. The effects of ERK5 on cancer onset and development have been widely reported [11, 12]. Our previous research indicated that the targeted blockage of the ERK5 pathway might be a latent treatment method against the development of BC [12]. In addition, mechanistic researches have investigated the involvement of the ERK5 pathway in oncocyte proliferation and cell cycle regulation, and these studies have found that such pathway is deregulated in several cancers [13]. Nevertheless, the functions of ERK5 in BC remain largely unknown.

Herein, we report that benzidine promoted cell growth and accelerated cell cycle transition, and it enhanced ERK5 activation in human BC cells UMUC-3. We found that decreased ERK5 via pharmacologic tools and genetic approaches reversed benzidine-induced proliferation. Collectively, the discoveries herein offer novel enlightenment regarding the function and mechanism of ERK5 in pathogenesis and provide a potential therapeutic target for benzidine-associated BC.

## MATERIALS AND METHODS

### Materials and reagents

Benzidine (4, 4’-diaminobiphenyl; ≥98.0%, RT), methyl alcohol, MTT and DMSO were provided by Merck (Reading Township, NJ, USA). Benzidine was dispersed in DMSO, which was afterwards stored under −20°C. The eventual level of DMSO was < 1‰. Rabbit anti-total ERK1/2 (no. 4376), rabbit anti-phosphorylated ERK1/2 (no. 4370), rabbit anti-p38 (no. 9212), rabbit anti-phosphorylated p38 (no. 9211), rabbit anti-JNK (no. 9252), rabbit anti-phosphorylated JNK (no. 4668), rabbit anti-total ERK5 (no. 12950), rabbit anti-phosphorylated ERK5 (no. 3371), rabbit anti-phosphorylated c-Jun (no. 2994), rabbit anti-phosphorylated c-Fos (no. 5348), rabbit anti-cyclin D1 (no. 55506), rabbit anti-PCNA (no. 13110), rabbit anti-p21 (no. 37543), and rabbit anti-GAPDH (no. 5174) antibodies were provided by CST (America). XMD8-92 was commercially obtained from Santa Cruz (America). Primers for cyclin D1, PCNA, p21, and GAPDH were produced by Invitrogen (America).

### Cellular cultivation and treatment

Human BC cell line UMUC-3 was provided by America Type Culture Collection (ATCC, America) and cultivated with DMEM added with 10% FBS (HyClone, America) in an incubating device with 5% CO_2_ under 37 °C. Cells were placed in 25-cm^2^ flasks, and the intermediary was changed every other day. When 80-90% confluency was reached, cells were exposed to different levels of benzidine for 5 days, with or without XMD8-92 or siRNAs specific to ERK5 (Invitrogen). Every procedure was completed for three times.

### Transfection

Prior to siRNA transfection, UMUC3 cells were cultivated in a 24-well dish. They were subjected to transfection via Lipofectamine 2000 (Invitrogen) as per standard protocols. The siRNA sequence for ERK5 was: 5’-GGGCCTATATCCAGAGCUU-3’. A scrambled control siRNA served as a negative control was acquired from Santa Cruz (America). The protein level of ERK5 was detected using western blot analysis at the fifth day after transfection.

### Cell viability assay

In short, cells were inoculated into 96-well dishes at 2.0×10^3^ cells/well and cultivated for one night with DMEM containing 10% FBS. To evaluate whether the suppression of ERK5 pathway might decrease benzidine-induced cell growth, three different sets of experiments were performed. (1) Cell growth analysis with benzidine alone: cells were exposed to different levels of benzidine (0-5 μM) for 5 days; (2) Cell growth analysis with a 5-day benzidine and XMD8-92 co-treatment: cells were treated with a specific content of benzidine (0.0075 μM) with/without XMD8-92 (5, 10, 15 μM) for 5 days; and (3) Cell growth analysis via a 5-day co-treatment of benzidine and siRNAs specific to ERK5: cells were treated with a specific content of benzidine (0.0075 μM) with/with no specific human siRNAs against ERK5 (10, 50, 100 pmol/L) for 5 days. At the end-point of assay, cells were treated with MTT solution for 4 hours. Posterior to the removal of MTT liquor and the solubilisation of crystals in DMSO, the OD was identified via a micro-plate reading device (Thermo Scientific, PRC) at a wave length of 490 nm.

### Cell cycle analysis

UMUC3 cells were inoculated into 6-well dishes at 1× 10^6^ cells/well, before the assay. They were then harvested, cleaned for two times in PBS and subjected to fixation in 70% ethyl alcohol under 4°C for one night. Posterior to the supplementation of 10 ml RNAse (10 mg/ml), cellular samples were left for 0.5 h under 37°C without light and dyed in 10 ml propidium iodide (1 mg/ml). Cells were subsequently analyzed via the FACScan Flow Cytometry Machine (BD Bioscience, America) to investigate the cellular cycle. The proportions of cells in every cellular cycle stage (G1, S, or G2) were computed via the MULTICYCLE 3.0 program (BD Bioscience), with a minimal 1× 10^5^ cells/ specimen assessed.

### WB analysis

UMUC3 cells were harvested, cleaned in cold PBS and subjected to lysis with RIPA buffering solution (Thermo Scientifific). The level of protein precipitation in cellular lysates was identified via the BCA Protein Analytical Tool (Pierce, America). Then, protein samples were subjected to 10% SDS-PAGE and moved to PVDF films (Millipore, America). After the blockade with 5% milk, the films were incubated with the first antibodies, and subsequently incubated with second antibodies. The detection was achieved using the chemiluminescent identification tool (Amersham Bioscience, America). The ImageJ program 1.4 (National Institution of Health) was utilized to study the gray level intensities of blot strips. The changes in band densities were presented by fold changes in contrast to the control in the blot posterior to the normalisation to GAPDH.

### qRT-PCR

Overall RNA was separated via RNAiso Plus as per the supplier’s protocols (TaKaRa, Japan). cDNA was produced from 2 mg overall RNA with AMV reversed transcriptive enzyme (TaKaRa) as per the supplier’s instruction. qRT-PCR was completed via the Power SYBR Green Master Mix (TaKaRa) and an ABI 7300 real-time PCR identification apparatus (Applied Biosystem, CA). All reactions were completed for three times. The cyclic profile was stated below: 95°C for 10 s, and 40 cycles of 95°C for 10 s and 72°C for 0.5 min. The comparative Ct approach was utilized for quantitative analysis, and the mRNA expression of each gene was normalised to GAPDH respectively. Every primer was produced by Invitrogen. The primers utilized are: GAPDH, forward 5’-GCTGCCCAACGCACCGAATA-3’ and reverse 5’-GAGTCAACGGATTTGGTCGT-3’; cyclin D1, forward 5’-CGTGGCCTCTAAGATGAAGG-3’ and reverse 5’-TGCGGATGATCTGTTTGTTC-3’; p21, forward 5’-GACACCACTGGAGGGTGACT-3’ and reverse 5’-CAGGTCCACATGGTCTTCCT-3’; PCNA, forward, 5’-TTTCACAAAAGCCACTCCACTG-3’; reverse, 5’-CTTTAAGTGTCCCATGTCAGCAAT-3’.

### Statistical analysis

Unless indicated otherwise, every assay was completed in triplicate. Data were described as mean ± SD posterior to the analysis via the SPSS 22.0 program (America). Unpaired Student’s t-test was utilized to compare 2 groups. One-way ANOVA was used for multi-group contrast, before the LSD significant difference test. Statistical significance was defined as p < 0.05.

## RESULTS

### Benzidine increased proliferative ability of UMUC3 cells

To examine the effects of benzidine on cellular activity, UMUC3 cells were exposed to benzidine at the concentrations from 0 to 5 μM. The outcomes herein revealed that 0.0025–0.01 μM benzidine exposure for 5 days remarkably elevated the cellular activity of UMUC3. Meanwhile, the treatment with benzidine at a concentration of 5 μM triggered cytotoxicity effects on UMUC3 cells (Fig. 1a). Consequently, certain benzidine levels (0.005 and 0.0075 μM) were selected for the subsequent procedures in our study.

**Fig. 1.**
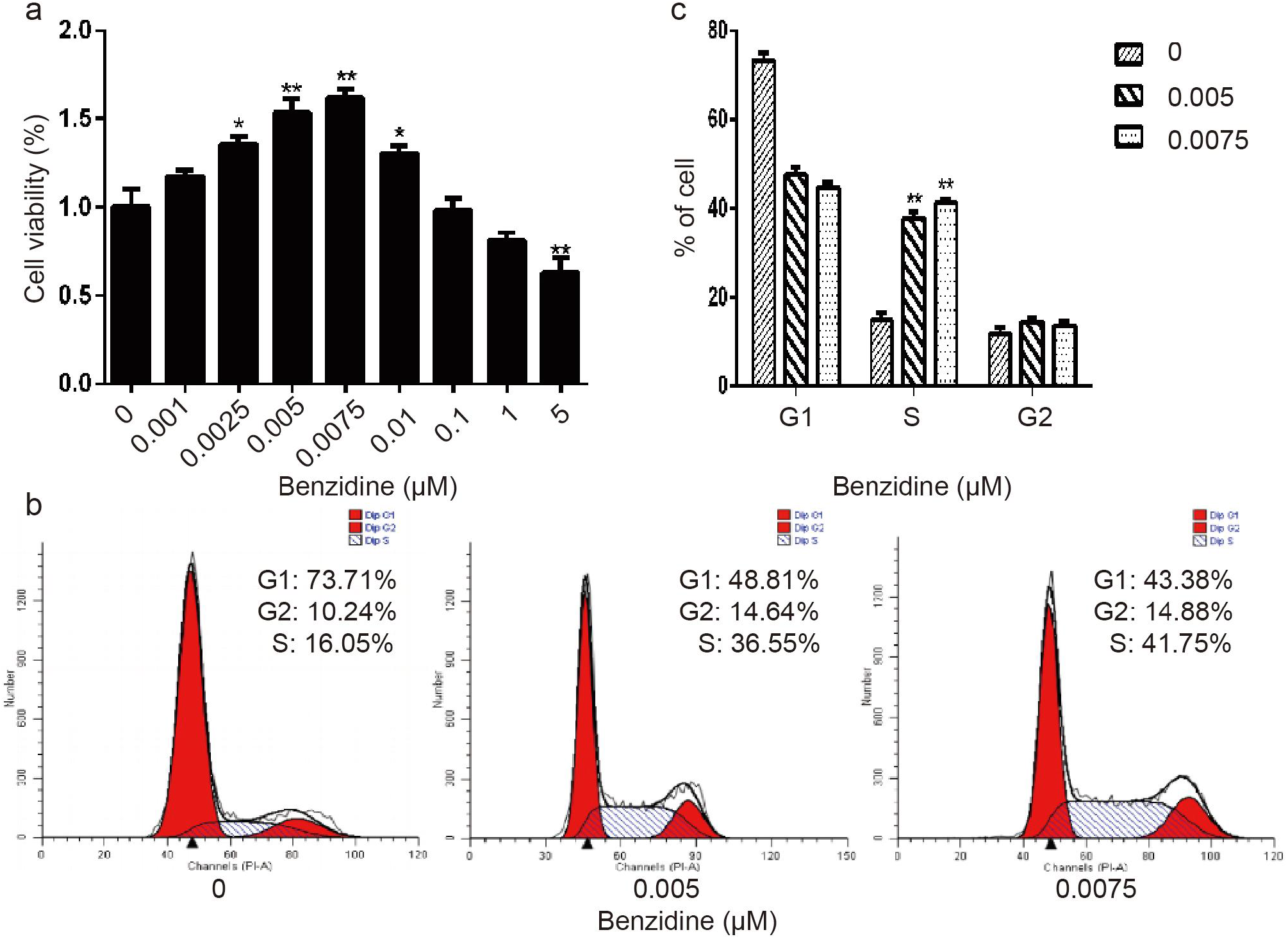
Benzidine induced the proliferative activity of human BC cells UMUC3. a) UMUC3 cells were exposed to diverse contents of benzidine for 5 days. Cellular activity was determined via MTT analysis. b and c) UMUC3 cells were exposed to indicated contents of benzidine (0, 0.005, 0.0075 μM) for 5 days. Then, the cellular cycle distribution was analyzed via flow cell technique. Data are described as average ± SD. *P < 0.05 and **P < 0.01, in contrast to the controls.

Flow cytometry outcomes unveiled that in contrast to the control cells, benzidine-treated UMUC3 cells had a remarkably influenced cellular cycle, where more cells shifted to the S stage and there were fewer G1 stage cells. This revealed that benzidine triggered G1/S transition and cellular proliferation (Fig. 1, b and c). Moreover, we studied the influence of benzidine on cell cycle via identifying the expressing levels of cell cycle regulatory proteins. We found that benzidine exposure enhanced the protein expressions of cyclin D1 and PCNA, whereas it decreased the contents of p21 (Fig. 2, a and b), which coincided with the results of qRT-PCR analysis (Fig. 2c). Collectively, those data demonstrated that benzidine exposure elicited UMUC3 cell proliferation.

**Fig. 2.**
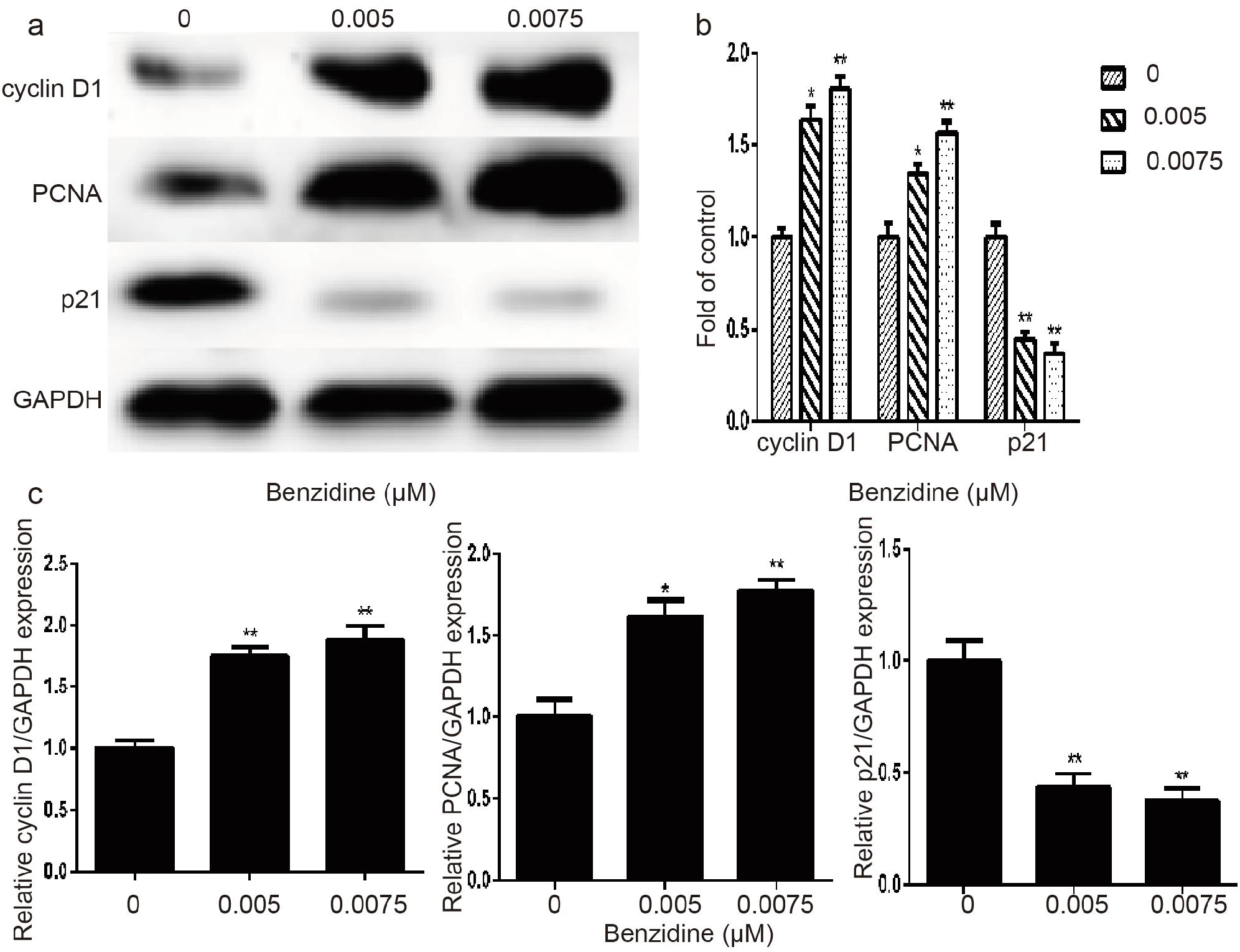
Benzidine changed the expressing levels of cell cycle genes in UMUC3 cells. a and b) Benzidine elevated the expression levels of cyclin D1 and PCNA proteins, and reduced the expression level of p21 protein, as displayed by WB analysis. GAPDH served as the load control. c) The mRNA levels of cyclin D1, PCNA, and p21 were determined by qRT-PCR, posterior to the normalisation to GAPDH. Data are presented as average ± SD. *P < 0.05 and **P < 0.01, in contrast to the controls.

### Benzidine activated ERK5/AP-1 pathway in UMUC3 cells

To evaluate the potential relevance between MAPK activation and benzidine-induced cell proliferation, the expressing levels of overall and phosphorylated ERK1/2, JNK, p38, ERK5 were identified by WB. We revealed that benzidine considerably stimulated the content of phosphorylated ERK5 in UMUC3 cells. We also discovered that benzidine had little impact on the expressing levels of p-ERK1/2, p-JNK, and p-p38 proteins (Fig. 3, a and b). In addition, benzidine treatment facilitated the stimulation of downstream target AP-1 proteins, as evidenced by the increased contents of phosphorylated c-Fos (p-c-Fos) and p-c-Jun (Fig. 3, c and d).

**Fig. 3.**
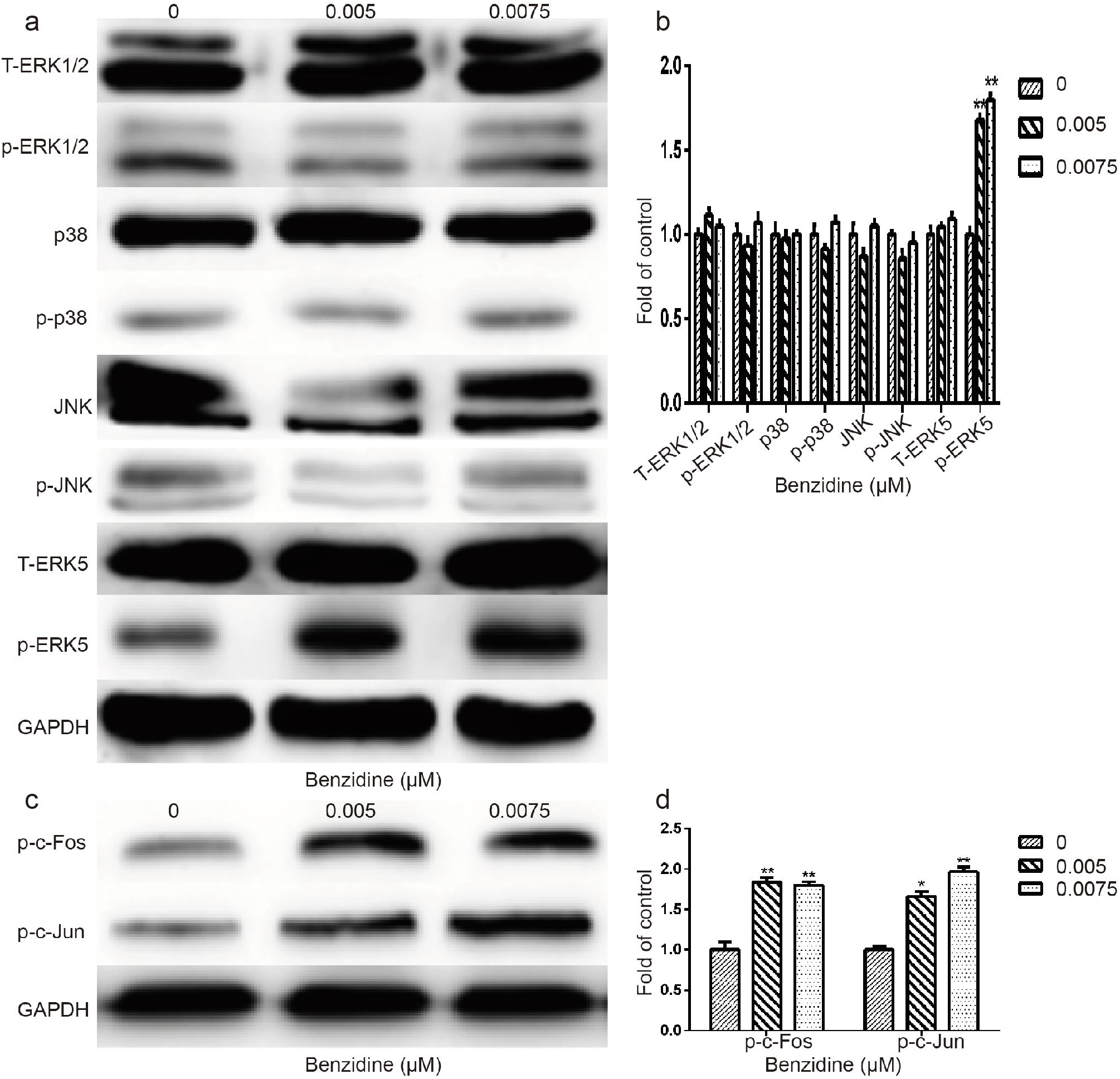
Benzidine-induced proliferation was associated with ERK5/AP-1 stimulation in UMUC3 cells. a) The expressing levels of T-ERK1/2, p-ERK1/2, p38, p-p38, JNK, p-JNK, T-ERK5 and p-ERK5 proteins via WB analysis. b) The expressing levels of AP-1 proteins (p-c-Fos and p-c-Jun) by WB analysis. GAPDH served as the load control. Data are presented as average ± SD. *P < 0.05 and **P < 0.01, in contrast to the controls.

### Blockage of ERK5 by XMD8-92 prevented benzidine-elicited UMUC3 cell proliferation

Subsequently, we explored if ERK5 facilitated benzidine-elicited cell proliferation in UMUC3. XMD8-92 (a specifific ERK5 inhibitor) was utilized to realize the pretreatment of cells. As presented by Fig. 4a, 10 μM XMD8-92 significantly rescued benzidine-triggered cell proliferation, as demonstrated by the cell viability assay. Moreover, flow cytometry unveiled that the fraction of cells in the S phase was remarkably reduced from 42.54% to 29.04% via XMD8-92 administration (Fig. 4, b and c). XMD8-92 reduced the expressing levels of p-ERK5, p-c-Fos, and p-c-Jun proteins in UMUC3 cells, as identified by WB (Fig. 5, a and b). In addition, based on WB analyses, XMD8-92 abolished the benzidine-activated decrease of p21 expressions, and it terminated the enhancement of cyclin D1 and PCNA (Fig. 5, c and d). Alike outcomes were discovered in qRT-PCR assay as well (Fig. 5e).

**Fig. 4.**
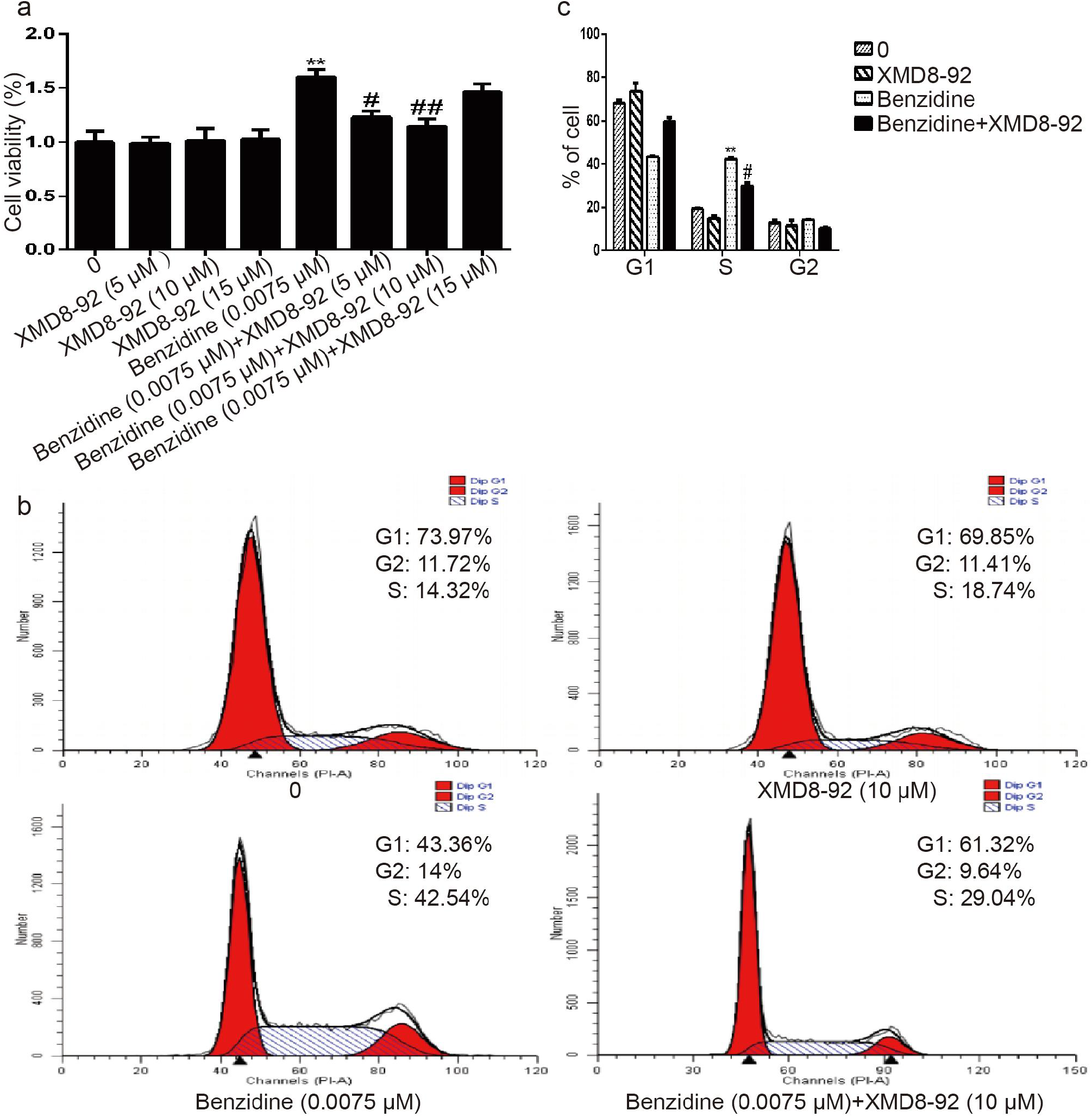
The suppression of ERK5 by XMD8-92 repressed benzidine-enhanced cellular proliferative activity and cellular cycle development in UMUC3 cells. a) Cells were exposed to diverse contents of XMD8-92 (0, 5, 10, 15 μM) with/with no 0.0075 μM benzidine for 5 days. Cellular activity was measured via MTT. **P < 0.01, in contrast to the controls; ^#^P < 0.05, ^##^P < 0.01, in contrast to the 0.0075 μM benzidine-exposed group. b and c) UMUC3 cells were exposed to 0.0075 μM benzidine with/with no 10 μM XMD8-92 for 5 days, and cellular cycle was analyzed via flow cytometry. Data are presented as average ± SD. **P < 0.01, in contrast to the controls; ^#^P < 0.05, in contrast to the 0.0075 μM benzidine-exposed group.

**Fig. 5.**
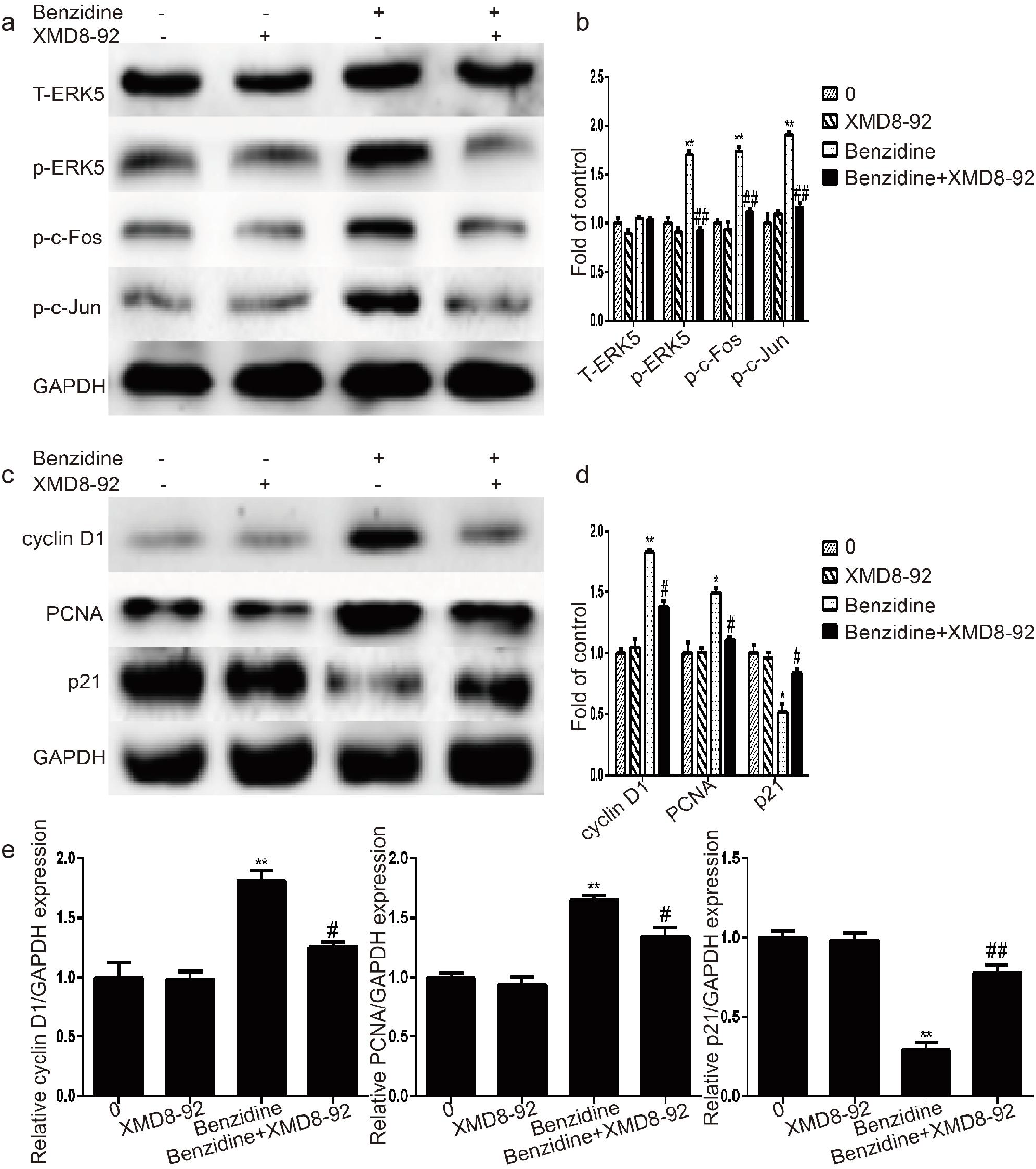
XMD8-92 reversed the benzidine-elicited variations of ERK5, AP-1 and cellular cycle proteins. UMUC3 cells were co-treated with 0.0075 μM benzidine and 10 μM XMD8-92 for 5 days. WB assay was utilized to identify the expressing levels of T-ERK5, p-ERK5, p-c-Jun, p-c-Fos (a and b), cyclin D1, PCNA, and p21 (c and d). GAPDH was the load control. XMD8-92 (10 μM) improved benzidine-elicited variations in the expressions of cyclin D1, PCNA, and p21 mRNAs e). Data are presented as average ± SD. *P < 0.05, **P < 0.01, in contrast to the controls; ^#^P < 0.05, ^##^P < 0.01, in contrast to the 0.0075 μM benzidine-exposed group.

### Inhibition of ERK5 by a specific siRNA reversed benzidine-enhanced UMUC3 cell growth

To further determine the link between ERK5 activation and benzidine-induced cell proliferation, we used ERK5-specific siRNA to silence ERK5 in UMUC3 cells. Initially, a MTT assay was performed to explore an appropriate transfection concentration of ERK5-specific siRNA. Results showed that transfection with 50 pmol/l siERK5 significantly decreased benzidine-enhanced cell growth. Meanwhile, 50 pmol/l scrambled control siRNA or siERK5 had little effect on the cell viability of UMUC3 (Fig. 6a). Flow cytometry revealed that siERK5 significantly decreased the S-phase population from 36.97 to 21.25% and increased the G1 fraction from 51.52 to 68.87%, which demonstrated that siERK5 restrained benzidine-elicited G1/S transition in UMUC3 cells (Fig. 6, b and c). As expected, WB analysis showed that siERK5 restored the benzidine-elicited elevation of p-ERK5, p-c-Fos, and p-c-Jun (Fig. 7, a and b). Moreover, siERK5 abolished benzidine-triggered changes in the expressing levels of cellular cycle modulatory proteins, as identified via WB (Fig. 7, c and d) and qRT-PCR (Fig. 7e). Collectively, those results revealed the pivotal effect of ERK5 on benzidine-induced UMUC3 cell proliferation.

**Fig. 6.**
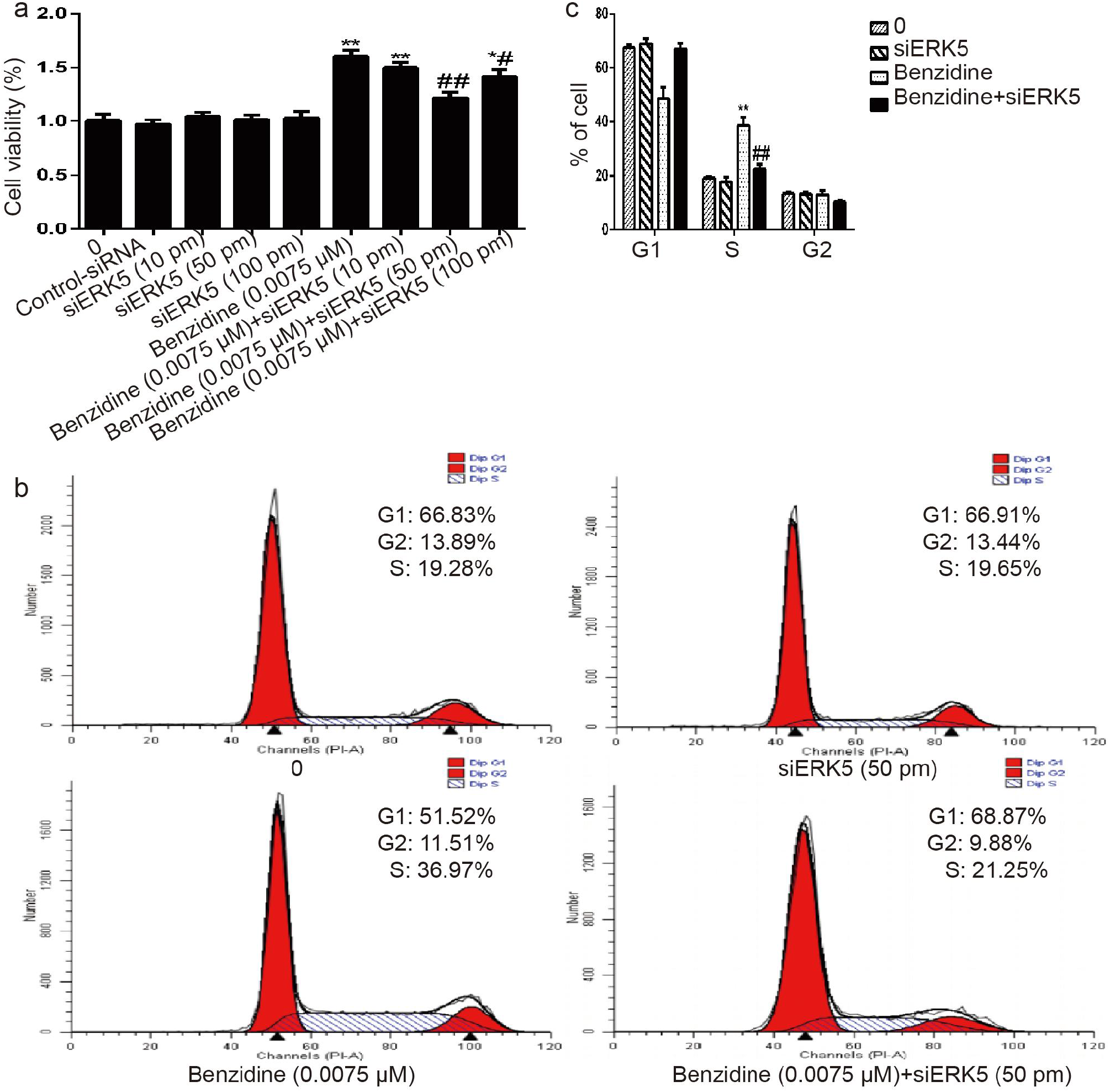
The suppression of ERK5 by siRNA attenuated benzidine-elicited proliferative activity and cellular cycle transition in UMUC3 cells. a) UMUC3 cells were transfected with indicated levels of ERK5-siRNA (10, 50, 100 pmol/l) and negative control (50 pmol/l). Then, cells were treated with/with no 0.0075 μM benzidine for 5 days. MTT analyses were completed to study cellular activity. b and c) Flow cytometry showed that benzidine-elicited cellular cycle transition was diminished by siRNA targeting ERK5. Data are presented as mean ± SD. *P < 0.05, **P < 0.01, in contrast to the controls; ^#^P < 0.05, ^##^P < 0.01, in contrast to the 0.0075 μM benzidine-exposed group.

**Fig. 7.**
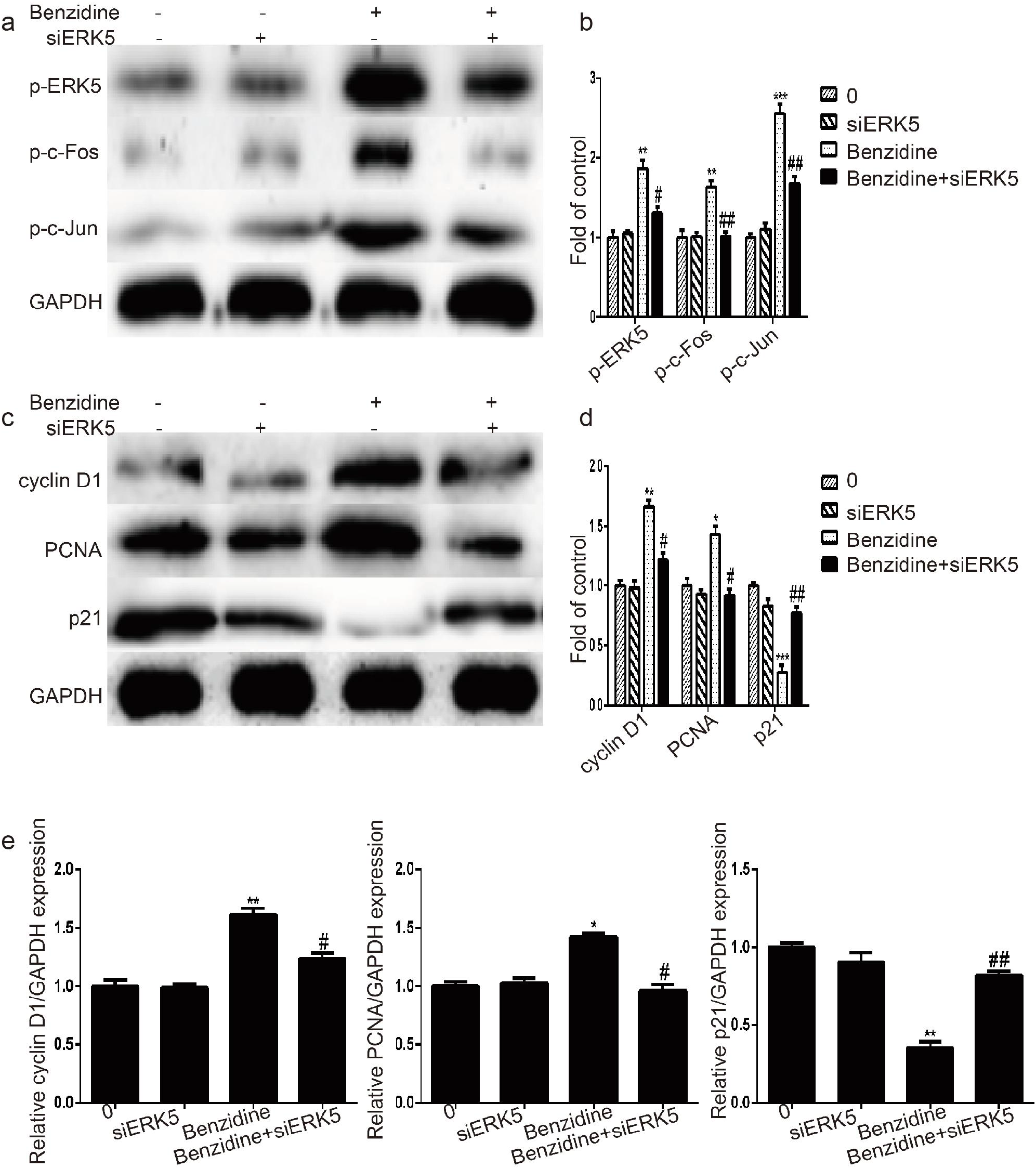
ERK5-specific siRNA prevented benzidine-triggered ERK5/AP-1 activation and variations in the expressions of cellular cycle proteins in UMUC3 cells. UMUC3 cells were transfected with 50 pmol/L siERK5 and 50 pmol/L non-silencing siRNA, or they were not transfected. Cells were subjected to lysis on the fifth day after transfection, and the protein levels of p-ERK5, p-c-Fos, p-c-Jun (a and b), cyclin D1, PCNA, and p21 (c and d) were identified via WB analysis. GAPDH served as the load control. e) The mRNA levels of cyclin D1, PCNA, and p21 based on qRT-PCR, after the normalisation to GAPDH. *P < 0.05, **P < 0.01, ***P < 0.001, in contrast to the controls; ^#^P < 0.05, ^##^P < 0.01, in contrast to the 0.0075 μM benzidine-exposed group.

## DISCUSSION

As the tenth most commonly seen cancer worldwide, BC has high morbidity and mortality rates. The lifetime risks of BC in males and females are approximately 1.1% and 0.27%, respectively [14]. Benzidine is a gene toxic substance and a crucial risk factor of bladder cancer. Substantial epidemiological researches concerning the association between benzidine and BC have been completed since 1950s [15]. In vitro and in vivo researches have assessed the genetic expression variations related to acute benzidine exposure [12, 16–22]. Previously, we reported that benzidine treatment could result in increased cellular proliferation, EMT alteration and MAPK activation in SV-HUC-1 and T24 cells [12, 17–21]. In addition, the enhancement of the stemness of BC stem cells was discovered after benzidine treatment [22]. Nonetheless, the causal links by which benzidine induced the developmental process of BC are still elusive. Herein, our team unveiled that benzidine treatment elicited cellular proliferative activities in human BC cells UMUC3. We unraveled that benzidine-enhanced cellular proliferation was related to the stimulation of the ERK5 path and AP-1 proteins. At the same time, the results herein revealed that the suppression of ERK5 by XMD8-92 or ERK5-specific siRNA avoided benzidine-elicited ERK5/AP-1 stimulation and abolished benzidine-elicited cellular proliferative activities in vitro.

There are three cell cycle checkpoints, namely G1/S, S and G2/M. Considered as the restriction point, the G1/S checkpoint is famous for its influence on ensuring abundant substances for DNA duplication and supervising DNA impairment [23]. G1 phase cells will accelerate the cellular cycle transition and transfer to the S phase when those cells are exposed to growth-facilitating stimulative substances induced by various kinds of chemicals like benzidine [17–19]. A few cellular molecules has been found to regulate the cell cycle of eukaryotic cells, which consist of cyclin D1, PCNA, and p21 [24–29]. Aberrant cellular cycle modulatory molecules have been identified in practically every mankind tumor type, including BC [7, 8, 17–19].

Cyclin D1 is a vital modulator of cellular cycle and is pivotal for the etiopathogenesis of tumor. As the allosteric modulator of the cyclin-dependent kinase (CDK) 4 and 6, cyclin D1 modulates the transition from G1 to S phase [24]. The overexpression of cyclin D1 drives unchecked cellular proliferation promoting multiple types of oncocyte growth, which include mammary carcinoma [25], hepatocellular carcinoma [26], and BC [8, 17–19], etc. Koppararu and colleagues showed that a higher cyclin D1 expression was found in 212 BC samples, in contrast to 131 neighboring healthy bladder samples. Different expression levels of cyclin D1 were discovered between muscle-invading BC and non-muscle-invading BC tissues, which suggested that cyclin D1 might be employed as a potential marker for determining the hypotypes of BC for precise treatment [27]. Intriguingly, sufferers with cyclin D1 positive cancers had a reduced survival rate in contrast to those with cyclin D1 negative cancers [28].

PCNA is a vital protein participating in various cellular processes, including DNA metabolic process, cell cycle control, cell death, and survival [29]. In normal human cells, PCNA is found to be related to a cyclin, a CDK and the CDK suppressor p21 to constitute a quaternary complex to avoid the G1-to-S transition [30]. Thereby, PCNA might be a molecule-level biomarker for cellular proliferative activities. The abnormal expression of PCNA is found to be related to BC progression. Skopelitou et al. indicated that PCNA expression was closely linked with the histological grade in BC and indirectly related to the tumour stage in BC [31]. The involvement of PCNA expression was also noted in various types of urinary cancer progression in vivo, including the bladder, prostate, and kidney [32–34]. Moreover, in vitro studies also discovered that high levels of PCNA were associated with benzidine-induced tumorigenesis and the development of BC [8, 17–19].

P21, also known as p21WAF1/Cip1, is a famous suppressor of cellular cycle and is capable of arresting the cellular cycle development in G1/S and G2/M transition. P21 can be elicited by p53-based and p53-based causal links. For the purpose of preserving genetic integrity, the p53-induced expression of p21 mediates G1 and G2 cell cycle arrest [35]. Interestingly, contradictory to its role as a suppressor, low levels of p21 facilitates proliferative activities through the stimulation of cyclin D/CDK 4 or CDK 6 complex in tumor lineage cells [36]. It is noteworthy that the downregulation of p21 expression promotes cell growth and cellular cycle development in reaction to various stimulating substances, such as cigarette smoke extract and benzidine [17–19, 37, 38]. In addition, p21 deregulation can also be noted in different types of cancer progression both in vivo and in vitro [39]. These researches indicate that p21 is not only a suppressor but a stimulator of the cellular cycle relying on the cell context and its expressing levels.

Previously, we reported that benzidine treatment elicited the cellular proliferative activities of T24 and SV-HUC-1 cells and triggered the G1-to-S transition, and it induced the expressing level alterations of cyclin D1, PCNA, and p21 [17–19]. In this study, we discovered that benzidine exposure enhanced UMUC3 cell proliferation, as demonstrated by the increased cell viability, the elevated proportion of cells in the S phase, and the varied expressing levels of cellular cycle regulatory proteins (like the decreased p21, as well as the increased cyclin D1 and PCNA).

Certain proofs have revealed that ERK5-mediated signal transduction facilitates the development of many tumors like gastric [40], bladder [12, 23], colon [41], and lung cancers [42]. ERK5 can be stimulated by receptor tyrosine kinases and other stimulating substances like benzidine [12, 17–22]. Mechanistic studies indicated that phosphorylated ERK5 transfered from the cytoplasm to the nucleus, and then regulated the activity of transcriptional factors to modulate cell proliferation [13]. The abnormal expression of ERK5 was detected in oncocytes, like stomach carcinoma MGC-803 cells, and it induced an enhanced ability of cell proliferation and colony formation, therefore contributing to increased cell growth [40]. Moreover, certain researches revealed that ERK5 regulated benzidine-mediated urocystic EMT and that ERK5 inhibition reversed EMT alterations [12, 22]. In a mouse model, fibroblast-specific ERK5 deficiency enhanced tumor vessel formation and increased the number of activated fibroblast, and it exacerbated tumor progression [41]. Sánchez-Fdez et al. indicated that the elevated contents of MEK5/ERK5 were related to unsatisfactory prognoses in pulmonary carcinoma. Subsequently the suppression of MEK5 or ERK5 by genetic and pharmacological tools restricted cancer development in vitro and in vivo [42]. The outcomes herein revealed that benzidine-elicited proliferative activities were related to ERK5/AP-1 stimulation in UMUC3 cells, whereas ERK1/2, p38, and JNK paths weren’t evidently implicated in such process. In addition, our team revealed that ERK5 suppression reversed benzidne-stimulated cell proliferation. Those results reveal that the development of BC cells along the cellular cycle needs the stimulation of the ERK5 pathway.

As a downstream target of the MAPK pathway and a critical regulator of transcription factor protein families, activator protein-1 (AP-1) is a group of proteins participating in various cellular processes [43]. AP-1 modulates the expressions and roles of cellular cycle modulators, like cyclin D1, PCNA, and p21, during cell proliferation [44, 45]. AP-1 overexpression is implicated in several kinds of mankind tumor specimens, like osteosarcoma [46], prostate carcinoma [47], etc. Notably, AP-1 was previously found to be associated with cancerogenesis and the development of benzidine-related BC [12, 17, 18, 20, 21]. In line with previous studies, we observed that benzidine-mediated ERK5 stimulation elevated the stimulation of AP-1 proteins. At the same time, benzidine-activated AP-1 stimulation was reduced posterior to the utilization of XMD8-92 or ERK5-siRNA. Those outcomes reveal the significance of AP-1 in benzidine-triggered cell proliferation of UMUC3.

To sum up, our research shows that benzidine enhances human BC UMUC3 cell proliferation via the stimulation of the ERK5/AP-1 pathway. These novel findings indicate that suppressing tumor growth by regulating ERK5 activity can provide new targets for benzidine-associated bladder cancer pharmacotherapies.

## Accepted abbreviations

BC: bladder cancer
MAPK: mitogen-activated protein kinase
ERK5: extracellular regulated protein kinases 5
MTT: 3-(4,5-dimethylthiazol-2-yl)-2,5-diphenyltetrazolium bromide
qRT-PCR: quantitative reverse transcription-polymerase chain reaction
siRNA: small interfering RNA
PCNA: proliferating cell nuclear antigen
AP-1: activator protein 1
ERK1/2: extracellular regulated protein kinases 1 and 2
JNKs: Jun N-terminal kinases
CDK: cyclin-dependent kinase.

## Acknowledgments

DY was responsible for the conception and design of the study. XS, TZ and LT acquired and analyzed the data. XS, TZ, LT, JM and YW were involved in drafting the manuscript, the interpretation of the data and revising the manuscript. All authors have read and approved the final manuscript.

## Ethics declarations

The authors declare no conflicts of interest in financial or any other sphere. This article does not contain descriptions of studies performed by the authors with participation of humans or using animals as object.

## REFERENCES

1. Cai, Z., and Liu, Q. (2021) Understanding the Global Cancer Statistics 2018: implications for cancer control, Sci. China Life Sci., 64(6), 1017–1020, doi: 10.1007/s11427-019-9816-1.

2. Siegel, R. L., Miller, K. D., Fuchs, H. E., and Jemal, A. (2022) Cancer statistics, 2022, CA. Cancer J. Clin., 72(1), 7–33, doi: 10.3322/caac.21708.

3. Millerick-May, M. L., Wang, L., Rice, C., and Rosenman, K. D. (2021) Ongoing risk of bladder cancer among former workers at the last benzidine manufacturing facility in the USA, Occup. Environ. Med., 78(9), 625–631, doi: 10.1136/oemed-2020-106431.

4. IARC (2020). IARC monographs on the evaluation of carcinogenic risks to humans, volume 100F-7. Benzidine, Lyon, France: International Agency for Research on Cancer, Available: https://monographs.iarc.fr/wp-content/uploads/2018/06/mono100F-7.pdf.

5. Chung, K. T. (2016) Azo dyes and human health: A review, J. Environ. Sci. Health C. Environ. Carcinog. Ecotoxicol. Rev., 34, 233–261, doi: 10.1080/10590501.2016.1236602.

6. Nurse, P. (2000) A long twentieth century of the cell cycle and beyond, Cell, 100, 71–78, doi: 10.1016/s0092-8674(00)81684-0.

7. Ingham, M., and Schwartz, G. K. (2017) Cell-Cycle Therapeutics Come of Age, J. Clin. Oncol., 35(25), 2949–2959, doi: 10.1200/JCO.2016.69.0032.

8. Zhao, F., Vakhrusheva, O., Markowitsch, S. D., Slade, K. S., Tsaur, I., Cinatl, J. J.r., Michaelis, M., Efferth, T., Haferkamp, A., and Juengel, E. (2020) Artesunate Impairs Growth in Cisplatin-Resistant Bladder Cancer Cells by Cell Cycle Arrest, Apoptosis and Autophagy Induction, Cells, 9(12), 2643, doi: 10.3390/cells9122643.

9. Iñesta-Vaquera, F. A., Campbell, D. G., Tournier, C., Gómez, N., Lizcano, J. M., and Cuenda, A. (2010) Alternative ERK5 regulation by phosphorylation during the cell cycle, Cell Signal, 22(12), 1829–1837, doi: 10.1016/j.cellsig.2010.07.010.

10. Drosten, M., and Barbacid, M. (2020) Targeting the MAPK Pathway in KRAS-Driven Tumors, Cancer Cell, 37(4), 543–550, doi: 10.1016/j.ccell.2020.03.013.

11. Monti, M., Celli, J., Missale, F., Cersosimo, F., Russo, M., Belloni, E., Di, M. A., Lonardi, S., Vermi, W., Ghigna, C., and Giurisato, E. (2022) Clinical Significance and Regulation of ERK5 Expression and Function in Cancer, Cancers (Basel), 14(2), 348, doi: 10.3390/cancers14020348.

12. Sun, X., Zhang, T., Deng, Q., Zhou, Q., Sun, X., Li, E., Yu, D., and Zhong, C. (2018) Benzidine Induces Epithelial-Mesenchymal Transition of Human Bladder Cancer Cells through Activation of ERK5 Pathway, Mol. Cells, 41(3), 188–197. doi: 10.14348/molcells.2018.2113.

13. Paudel, R., Fusi, L., and Schmidt, M. (2021) The MEK5/ERK5 pathway in health and disease, Int. J. Mol. Sci., 22, 7594, doi: 10.3390/ijms22147594.

14. Cumberbatch, M. G. K., Jubber, I., Black, P. C., Esperto, F., Figueroa, J. D., Kamat, A. M., Kiemeney, L., Lotan, Y., Pang, K., Silverman, D. T., Znaor, A., and Catto, J. W. F. (2018) Epidemiology of bladder cancer: a systematic review and contemporary update of risk factors in 2018, Eur. Urol., 74(6), 784–795, doi: 10.1016/j.eururo.2018.09.001.

15. Kolli, R. T., Xu, Z. L., Panduri, V., and Taylor J. A. (2021) Differential Gene Expression in Bladder Tumors from Workers Occupationally Exposed to Arylamines, Biomed. Res. Int., 2021, 2624433, doi: 10.1155/2021/2624433.

16. Chen, S. C., Hseu, Y. C., Sung, J. C., Chen, C. H., Chen, L. C., and Chung, K. T. (2011) Induction of DNA damage signaling genes in benzidine-treated HepG2 cells, Environ. Mol. Mutagen., 52(8), 664–672, doi: 10.1002/em.20669.

17. Sun, X., Deng, Q., Liang, Z., Zhang, Z., Zhao, L., Geng, H., Xie, D., Wang, Y., Yu, D., and Zhong, C. (2016) Curcumin reverses benzidine-induced cell proliferation by suppressing ERK1/2 pathway in human bladder cancer T24 cells, Exp. Toxicol. Pathol., 68(4), 215–222, doi: 10.1016/j.etp.2015.12.003.

18. Zhao, L., Zhang, T., Geng, H., Liu, Z., Liang, Z., Zhang, Z., Min, J., Yu, D., and Zhong, C. (2018) MAPK/AP-1 pathway regulates benzidine-induced cell proliferation through the control of cell cycle in human normal bladder epithelial cells, Oncol. Lett., 16(4), 4628–4634, doi: 10.3892/ol.2018.9155.

19. Ding, D., Liu, Z., Zhao, L., Geng, H., Liang, Z., and Yu, D. (2019) Role of PI3K/Akt pathway in Benzidine-induced proliferation in SV-40 immortalized human uroepithelial cell, Transl. Cancer Res., 8(4), 1301–1310. doi: 10.21037/tcr.2019.07.14.

20. Zhao, L., Geng, H., Liang, Z., Zhang, Z., Zhang, T., Yu, D., and Zhong, C. (2015) Benzidine induces epithelial-mesenchymal transition in human uroepithelial cells through ERK1/2 pathway, Biochem. Biophys. Res. Commun., 459(4), 643–649, doi: 10.1016/j.bbrc.2015.02.163.

21. Liu, Z., Liu, J., Zhao, L., Geng, H., Ma, J., Zhang, Z., Yu, D., and Zhong, C. (2017) Curcumin reverses benzidine-induced epithelial-mesenchymal transition via suppression of ERK5/AP-1 in SV-40 immortalized human urothelial cells, Int. J. Oncol., 50(4), 1321–1329, doi: 10.3892/ijo.2017.3887.

22. Wang, D., Xie, D., Bi, L., Wang, Y., Zou, C., Chen, L., Geng, H., Qian, W., Li, Y., Sun, H., Wang, X., Lu, Y., Yu, D., and Zhong, C. (2021) Benzidine promotes the stemness of bladder cancer stem cells via activation of the Sonic hedgehog pathway, Oncol. Lett., 21(2), 146, doi: 10.3892/ol.2020.12407.

23. Fu, G. H., Xu, Z. J., Chen, X. Y., Pan, H., Wang, Y. M., and Jin B. Y. (2020) CDCA5 functions as a tumor promoter in bladder cancer by dysregulating mitochondria-mediated apoptosis, cell cycle regulation and PI3k/AKT/mTOR pathway activation, J. Cancer, 11(9), 2408–2420, doi: 10.7150/jca.35372.

24. Montalto, F. I., and Amicis, F. D. (2020) Cyclin D1 in Cancer: A Molecular Connection for Cell Cycle Control, Adhesion and Invasion in Tumor and Stroma, Cells, 9(12), 2648, doi: 10.3390/cells9122648.

25. Komini, C., Theohari, I., Lambrianidou, A., Nakopoulou, L., and Trangas, T. (2021) PAPOLA contributes to cyclin D1 mRNA alternative polyadenylation and promotes breast cancer cell proliferation, J. Cell Sci., 134(7), jcs252304, doi: 10.1242/jcs.252304.

26. Wang, M., and Zhao, H. (2020) LncRNA CTBP1-AS2 promotes Cell Proliferation in Hepatocellular Carcinoma by Regulating the miR-623/Cyclin D1 Axis, Cancer Biother. Radiopharm., 35(10), 765–770, doi: 10.1089/cbr.2019.3375.

27. Kopparapu, P. K., Boorjian, S. A., Robinson, B. D., Downes, M., Gudas, L. J., Mongan, N. P., and Persson, J. L. (2013) Expression of cyclin d1 and its association with disease characteristics in bladder cancer, Anticancer Res., 33(12), 5235–5242.

28. Moradi, B. M., Bahrami, A., Khazaei, M., Ryzhikov, M., Ferns, G. A., Avan, A., and Mahdi, H. S. (2020) The prognostic value of cyclin D1 expression in the survival of cancer patients: A meta-analysis, Gene, 728, 144283, doi: 10.1016/j.gene.2019.144283.

29. Cardano, M., Tribioli, C., and Prosperi, E. (2020) Targeting Proliferating Cell Nuclear Antigen (PCNA) as an Effective Strategy to Inhibit Tumor Cell Proliferation, Curr. Cancer Drug Targets, 20(4), 240–252, doi: 10.2174/1568009620666200115162814.

30. He, G., Kuang, J., Koomen, J., Kobayashi, R., Khokhar, A. R., and Siddik, Z. H. (2013) Recruitment of trimeric proliferating cell nuclear antigen by G1-phase cyclin-dependent kinases following DNA damage with platinum-based antitumour agents, Br. J. Cancer, 109(9), 2378–2388, doi: 10.1038/bjc.2013.613.

31. Skopelitou, A., Korkolopoulou, P., Papanicolaou A, Christodoulou, P., Thomas-Tsagli, E., and Pavlakis, K. (1991) Comparative assessment of proliferating cell nuclear antigen immunostaining and of nucleolar organizer region staining in transitional cell carcinomas of the urinary bladder, Eur. Urol., 22, 235–240, doi: 10.1159/000474762.

32. Zuo, Y., Xu, X., Chen, M., and Qi, L. (2021) The oncogenic role of the cerebral endothelial cell adhesion molecule (CERCAM) in bladder cancer cells in vitro and in vivo, Cancer Med., 10(13), 4437–4450, doi: 10.1002/cam4.3955.

33. Chen, C. J., Cai, Z. D., Zhuo, Y. J., Xi, M., Lin, Z. Y., Jiang, F. N., Liu, Z. Z., Wan, Y. P., Zheng, Y., Li, J. X., Zhou, X., Zhu, J. G., Zhong, W. D. (2020) Overexpression of SLC6A1 associates with drug resistance and poor prognosis in prostate cancer, BMC Cancer, 20(1), 289, doi: 10.1186/s12885-020-06776-7.

34. Yang, C. M., Ji, S., Li, Y., Fu, L. Y., Jiang, T., and Meng, F. D. (2017) β-Catenin promotes cell proliferation, migration, and invasion but induces apoptosis in renal cell carcinoma, Onco. Targets Ther., 10, 711–724, doi: 10.2147/OTT.S117933.

35. Bogdanow, B., Phan, Q. V., and Wiebusch, L. (2021) Emerging Mechanisms of G 1/S Cell Cycle Control by Human and Mouse Cytomegaloviruses, mBio, 12(6), e0293421, doi: 10.1128/mBio.02934-21.

36. Cheng, M., Olivier, P., Diehl, J. A., Fero, M., Roussel, M. F., Roberts, J. M., and Sherr, C. J. (1999) The p21(Cip1) and p27(Kip1) CDK ‘inhibitors’ are essential activators of cyclin D-dependent kinases in murine fifibroblasts, EMBO J., 18, 1571–1583, doi: 10.1093/emboj/18.6.1571.

37. Geng, H., Zhao, L., Liang, Z., Zhang, Z., Xie, D., Bi, L., Wang, Y., Zhang, T., Cheng, L., Yu, D., and Zhong, C. (2017) Cigarette smoke extract-induced proliferation of normal human urothelial cells via the MAPK/AP-1 pathway, Oncol. Lett., 13(1), 469–475, doi: 10.3892/ol.2016.5407.

38. Deng, Q., Sun, X., Liang, Z., Zhang, Z., Yu, D., and Zhong C. (2016) Cigarette smoke extract induces the proliferation of normal human urothelial cells through the NF-κB pathway, Oncol. Rep., 35(5), 2665–2672, doi: 10.3892/or.2016.4623.

39. Kreis, N. N., Louwen, F., and Yuan, J. P. (2019) The Multifaceted p21 (Cip1/Waf1/ CDKN1A) in Cell Differentiation, Migration and Cancer Therapy, Cancers (Basel), 11(9), 1220, doi: 10.3390/cancers11091220.

40. Wu, L. C., Chen, J. S., Qin, Y. Z., Mo, X. W., Huang, M. W., Ru, H. M., Yang, Y., Liu, J. G., and Lin, Y. (2016) SATB2 suppresses gastric cancer cell proliferation and migration, Tumour Biol., 37(4), 4597–4602, doi: 10.1007/s13277-015-4282-5.

41. Imanishi, M., Yamakawa, Y., Fukushima, K., Ikuto, R., Maegawa, A., Izawa-Ishizawa, Y., Horinouchi, Y., Kondo, M., Kishuku, M., Goda, M., Zamami, Y., Takechi, K., Chuma, M., Ikeda, Y., Tsuchiya, K., Fujino, H., Tsuneyama, K., and Ishizawa, K. (2020) Fibroblast-specific ERK5 deficiency changes tumor vasculature and exacerbates tumor progression in a mouse model, Naunyn Schmiedebergs Arch. Pharmacol., 393(7), 1239–1250, doi: 10.1007/s00210-020-01859-5.

42. Sánchez-Fdez, A., Re-Louhau, M. F., Rodríguez-Núñez, P., Ludeña, D., Matilla-Almazán, S., Pandiella, A., and Esparís-Ogando, A. (2021) Clinical, genetic and pharmacological data support targeting the MEK5/ERK5 module in lung cancer, NPJ Precis. Oncol., 5(1), 78, doi: 10.1038/s41698-021-00218-8.

43. Atsaves, V., Leventaki, V., Rassidakis, G. Z., and Claret, F. X. (2019) AP-1 Transcription Factors as Regulators of Immune Responses in Cancer, Cancers (Basel), 11(7), 1037, doi: 10.3390/cancers11071037.

44. Shaulian, E., and Karin, M. (2001) AP-1 in cell proliferation and survival, Oncogene, 20(19), 2390–400, doi: 10.1016/s0955-0674(97)80068-3.

45. Liu, Y. C., Chang, H. W., Lai, Y. C., Ding, S. T., and Ho, J. L. (1998) Serum responsiveness of the rat PCNA promoter involves the proximal ATF and AP-1 sites, FEBS Lett., 441(2), 200–204, doi: 10.1016/s0014-5793(98)01549-x.

46. Matsuoka, K., Bakiri, L., Wolff, L. I., Linder, M., Mikels-Vigdal, A., Patiño-García, A., Lecanda, F., Hartmann, C., Sibilia, M., and Wagner, E. F. (2020) Wnt signaling and Loxl2 promote aggressive osteosarcoma, Cell Res., 30(10), 885–901, doi: 10.1038/s41422-020-0370-1.

47. Zhang, S., Rong, P., Chen, Q., and Wang, W. (2019) Suppressor of activator protein-1 regulated by interferon expression in prostate cancer tissues and cells, Life Sci., 232, 116626, doi: 10.1016/j.lfs.2019.116626.

